# The molecular chaperone Cysteine String Protein is required to stabilize *trans*-SNARE complexes during human sperm acrosomal exocytosis

**DOI:** 10.1101/2022.03.03.482836

**Authors:** Karina Flores Montero, María Victoria Berberián, Luis Segundo Mayorga, Claudia Nora Tomes, María Celeste Ruete

## Abstract

Membrane fusion in sperm cells is crucial for acrosomal exocytosis and must be preserved to assure fertilizing capacity. Evolutionarily conserved protein machinery regulates acrosomal exocytosis. Molecular chaperones play a vital role in spermatogenesis and post-testicular maturation. Cysteine String Protein (CSP) is a member of the Hsp40 co-chaperones, and for more than 20 years, most research published focused on CSP’s role in synapsis. However, the participation of molecular chaperones in acrosomal exocytosis is poorly understood. Using western blot and indirect immunofluorescence, we showed that CSP is present in human sperm, is predominantly bound to membranes, and is palmitoylated. Moreover, using electron microscopy and functional assays, we reported that sequestration of CSP avoided the assembly of *trans*-complexes and inhibited exocytosis. In summary, our data demonstrated that CSP is necessary and mediates the *trans*-SNARE complex assembly between the outer acrosomal and plasma membranes, thereby regulating human sperm acrosomal exocytosis. Understanding CSP’s role is critical in identifying new biomarkers and generating new rational-based approaches to treating male infertility.

**Summary statement:** Cysteine String Protein is necessary and mediates the *trans*-SNARE complex assembly between the outer acrosomal and plasma membranes, thereby regulating human sperm acrosomal exocytosis.

## INTRODUCTION

Male infertility is a complex disorder; about 10-20% is idiopathic (Leslie *et al*., 2021), and the underlying mechanisms are not entirely understood. Several molecular chaperones play a critical role in spermatogenesis and post-testicular maturation (Dun, Aitken and Nixon, 2012). Evidence suggests that anomalous chaperone expression may be a determinant factor leading to male infertility. However, far too little attention has been directed to the molecular chaperone Cysteine String Protein in acrosomal exocytosis.

The presence of a protein-folding machinery is crucial for the proper folding of proteins. Molecular chaperones comprise a family of proteins that interact with hydrophobic domains exposed transiently in their targets. Chaperones drive the correct folding in native conformation of nascent polypeptides (Hartl, Bracher and Hayer-Hartl, 2011). Also, they participate in unfolding or in preventing inappropriate aggregation to ensure a productive folding of the proteins themselves, or the protein-complex association (Dun, Aitken and Nixon, 2012; Gorenberg and Chandra, 2017). Cysteine String Proteins (CSP) are members of the Hsp40/DNAJ co-chaperones that contribute to exocytosis (Gundersen, Mastrogiacomo and Umbach, 1995). According to subcellular location and tissue distribution, these members have been divided into three subtypes; among these, CSP belongs to subtype III (DnaJC) (Cheetham and Caplan, 1998). The structure of CSP is subdivided into four domains: (i) a conserved J-domain near the amino terminus responsible for the HSP70 interaction and ATPase activity regulation (Minami *et al*., 1996; Cheetham and Caplan, 1998); (ii) a linker domain that joins the J-domain and cysteine-string portion (Boal *et al*., 2011); (iii) a cysteine-rich string domain that contains 14 cysteines palmitoylated in vivo (Gundersen *et al*., 1994), this palmitoylation may be essential to maintain the proper membrane orientation of CSP (Greaves and Chamberlain, 2006); and (iv) a less characterized C-terminal portion. Specifically, CSPα (DnaJC5α) localizes to synaptic vesicles playing a role in exocytosis in the nervous system (Chandra *et al*., 2005), and promotes SNARE-complex assembly during synaptic activity (Sharma, Burré and Südhof, 2011). Also, it is implicated in exosomes (Deng *et al*., 2017), and other protein secretion pathways like release of neurodegenerative disease proteins (Fontaine *et al*., 2016; Lee *et al*., 2016). Moreover, CSPβ (DnaJC5β) isoform is preferentially expressed in testis (Gorleku and Chamberlain, 2010) and associated with nerve terminals in the mouse brain (Gundersen *et al*., 2010).

CSPα has a well-characterized role in regulated exocytosis at nerve terminals (Gundersen, 2020). Membrane fusion is executed in a specific moment by conserved protein machinery during the evolution of the eukaryotic cells (Südhof and Rizo, 2011). In sperm cells, membrane fusion is a critical step in the process of acrosomal exocytosis, is regulated by calcium signaling and must be preserved to assure fertilizing capacity. The acrosome content assist sperm penetration and facilitate disruption of the egg coat and, finally, oocyte fertilization (Hirohashi and Yanagimachi, 2018).

The central components of the eukaryotic fusion machinery in synaptic transmission are the SNAREs (soluble, N-ethylmaleimide-sensitive attachment receptors) synaptobrevin-2, syntaxin-1, and SNAP-25 (synaptosome-associated protein of 25 kDa) (Jahn and Scheller, 2006; Südhof and Rizo, 2011). These proteins are localized in the outer acrosomal membrane and the sperm plasma membrane and form a highly stable four-helix bundle called SNARE complex (Tomes, 2015). In sperm cells, *the cis-*SNARE complex is present until the trigger of the acrosome exocytosis. Consequently, SNAREs become monomeric. Finally, acrosome docking is conducted by transitioning to a productive *trans*-SNARE complex assembly (Tomes *et al*., 2002; De_Blas *et al*., 2005). SNARE complex assembly is topologically complex and is exquisitely controlled by key proteins such as Munc13, Munc18, synaptotagmin and complexin (Roggero *et al*., 2007; Bello *et al*., 2012; Rodríguez *et al*., 2012; Tsai *et al*., 2012; Prinslow *et al*., 2019).

To date there are several observations concerning the molecular role of CSPs in regulated exocytosis at different cell types. CSPα knockout causes synapse loss and neurodegeneration in mice (Fernández-Chacón *et al*., 2004; Donnelier and Braun, 2014; Lopez-Ortega, Ruiz and Tabares, 2017; Valenzuela-Villatoro *et al*., 2018). While some research has been conducted on CSPα’s role, very little is known about the importance of CSPβ in regulated exocytosis. Notwithstanding studies that revealed that this protein has a higher expression in testis, CSPβ is poorly studied (Boal *et al*., 2007).

A better comprehension of the molecular mediators connecting *trans*-SNARE assembly and acrosomal exocytosis may reveal clinically relevant pathways leading to infertility. Knowing the mechanisms of chaperone actions in the sperm and their possible participation in infertility would allow the development of therapeutic-targeting strategies to solve unexplained male fertility problems. The involvement of CSP in the regulation of acrosomal exocytosis has not been reported. Therefore, the main goal here was to evaluate the role of CSP in the mechanism of human sperm acrosomal exocytosis.

## RESULTS

### CSP is present in the acrosomal region of human sperm

The Human Protein Atlas (proteinatlas.org) (Uhlén *et al*., 2015) provides a complete picture of protein expression profiles in diverse human normal tissues. The consensus of three databases shows fifty-five different tissue and cell types with high-medium CSPα mRNA expression (Fig. 1A). However, DnaJC5β expression was shown to be almost exclusively expressed in testis (Fig. 1B). Interestingly, single-cell RNAseq cluster analysis showed cell-type specificity of CSPβ in late and early spermatids (Fig. 1C). Altogether these data revealed that CSPα appears to be associated with neural tissues, and CSPβ is predominantly present in testis. In this regard, CSPβ RNA expression is significantly higher in testis compared to CSPα (Fig. 1D).

**Figure 1.**
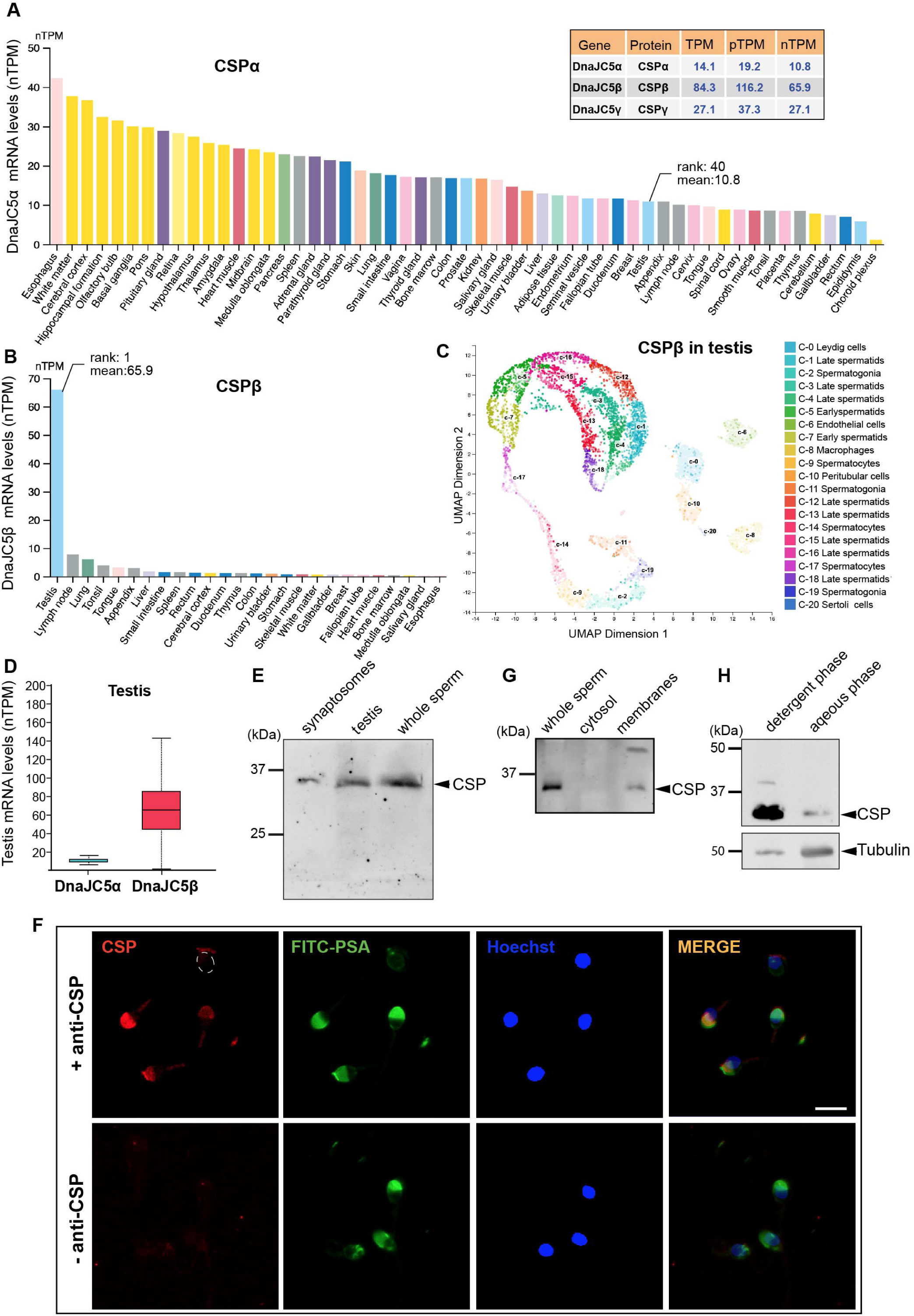
CSP expression profiles in human tissues and protein present in human sperm and located in the acrosomal region. (A-B) The mRNA expression profiles (“nPTM”: normalized transcripts per million) of DnaJC5α (**A**) and DnaJC5β (**B**) in human tissues, created by combining the data from the three transcriptomics datasets (HPA, GTEx, and FANTOM5) (https://www.proteinatlas.org/ENSG00000147570-DNAJC5B/tissue). (**C**) Uniform Manifold Approximation and Projection (UMAP) clustering of single cell data representing the identified cellular clusters that are colored by cell type (https://www.proteinatlas.org/ENSG00000147570-DNAJC5B/single+cell+type/testis). (**D**) Comparison of CSPα (Average nTPM: 10.8) and CSPβ (Average nTPM: 65.9) mRNA expression in normal testis (https://www.proteinatlas.org/ENSG00000147570-DNAJC5B/tissue/testis). (**E**) Proteins from whole human sperm extract, mouse testis, and synaptosomes were electrophoresed in 10% Tris-glycine-SDS-PAGE, transferred to a nitrocellulose membrane, and immunoblotted with an antibody raised against CSP as described in Material and Methods. The molecular mass standards are indicated to the left. Black arrows point to the 34 kDa specific CSP bands. (**G**) Sperm homogenates (100 × 10^6^ cells) were subjected to subcellular fractionation into soluble (cytosol) and particulate (membranes) fractions were performed as described in Material and Methods. Then were electrophoresed in 10% Tris-glycine-SDS-PAGE, transferred to a nitrocellulose membrane, and immunoblotted with anti-CSP (1 µg/ml). (**H**) Whole-sperm homogenate (200 × 10^6^ cells) was partitioned in Triton X-114. Cell partition in an aqueous and a detergent phase was done as described in Material and Methods. Then, the phases were electrophoresed in 10% Tris-glycine-SDS-PAGE, transferred to a nitrocellulose membrane, and immunoblotted with anti-CSP (1 µg/ml). (**F**) Capacitated human sperm were immunostained with (+anti-CSP) or without (-anti-CSP) antibody to CSP (1 µg/ml). Cells were triple stained with a fluorescent anti-rabbit antibody as a read-out for the presence of CSP (anti-rabbit-Cy3, red), FITC-PSA (green) to visualize intact acrosomes, and nuclei were stained with Hoechst 33342 (blue). The contour of the head is marked with a dotted oval (top and bottom). Scale bar = 5 µm. Shown are representative images of three independent experiments.

As we mentioned, there is no evidence of the implication of CSP in the mechanism of exocytosis in human sperm. To investigate the presence of CSP in human sperm, we prepared extracts from the whole human capacitated sperm and resolved it in 10% SDS-PAGE. We detected a band with a molecular mass of 34 kDa corresponding to CSP protein (Fig. 1E). We used mouse brain synaptosomes and human testis as positive controls and the antibody against CSP recognized bands in both human and mouse samples.

Next, we moved forward to evaluate the CSP distribution in human sperm. Indirect immunofluorescence microscopy revealed CSP localization entirely in the acrosome region (Fig. 1F, top). Also, note that the CSP mark disappeared when the acrosome was lost (Fig. 1F, oval contour, top). CSPs contain multiple cysteines within their cysteine-string domain, most of which are palmitoylated (Gundersen *et al*., 1994). Considering this posttranslational modification participates in the CSP membrane association (Greaves and Chamberlain, 2006), we decided to investigate the subcellular distribution of CSP in human sperm. To define the precise localization of CSP within cellular compartments we performed subcellular fractionation of human sperm into cytosolic and membrane fractions. All CSP was visualized in the membrane-associated fraction (Fig. 1G). This evidence agreed with our observation of the CSP molecular weight for a fully palmitoylated protein (Fig. 1E). Additionally, we phase-separated sperm in Triton X-114 detergent and evaluated partitions in the aqueous (cytosolic fraction) and detergent (cell membrane fraction) phase. Data showed that CSP predominantly was extracted in the detergent phase confirming its association with the membranes (Fig. 1H).

### CSP is required for acrosomal exocytosis

The presence of CSP in the acrosomal membrane region of sperm suggested that this might have a role in acrosomal exocytosis. To test this hypothesis, we sequestered endogenous CSP incubating with a specific antibody and analysed the outcome in acrosomal. We incubated SLO permeabilized human sperm with increasing concentrations of anti-CSP antibody and evaluated its effect on calcium-stimulated acrosome reaction. As shown in Fig. 2A, the antibody inhibited exocytosis in a dose-dependent manner. Then, we treated permeabilized sperm with anti-CSP, stimulated the acrosomal exocytosis with calcium, and added a recombinant non-palmitoylated CSPβ. Interestingly, the addition of the exogenous CSPβ reversed the blockade imposed by the antibody (Fig. 2B, top yellow bar).

**Figure 2.**
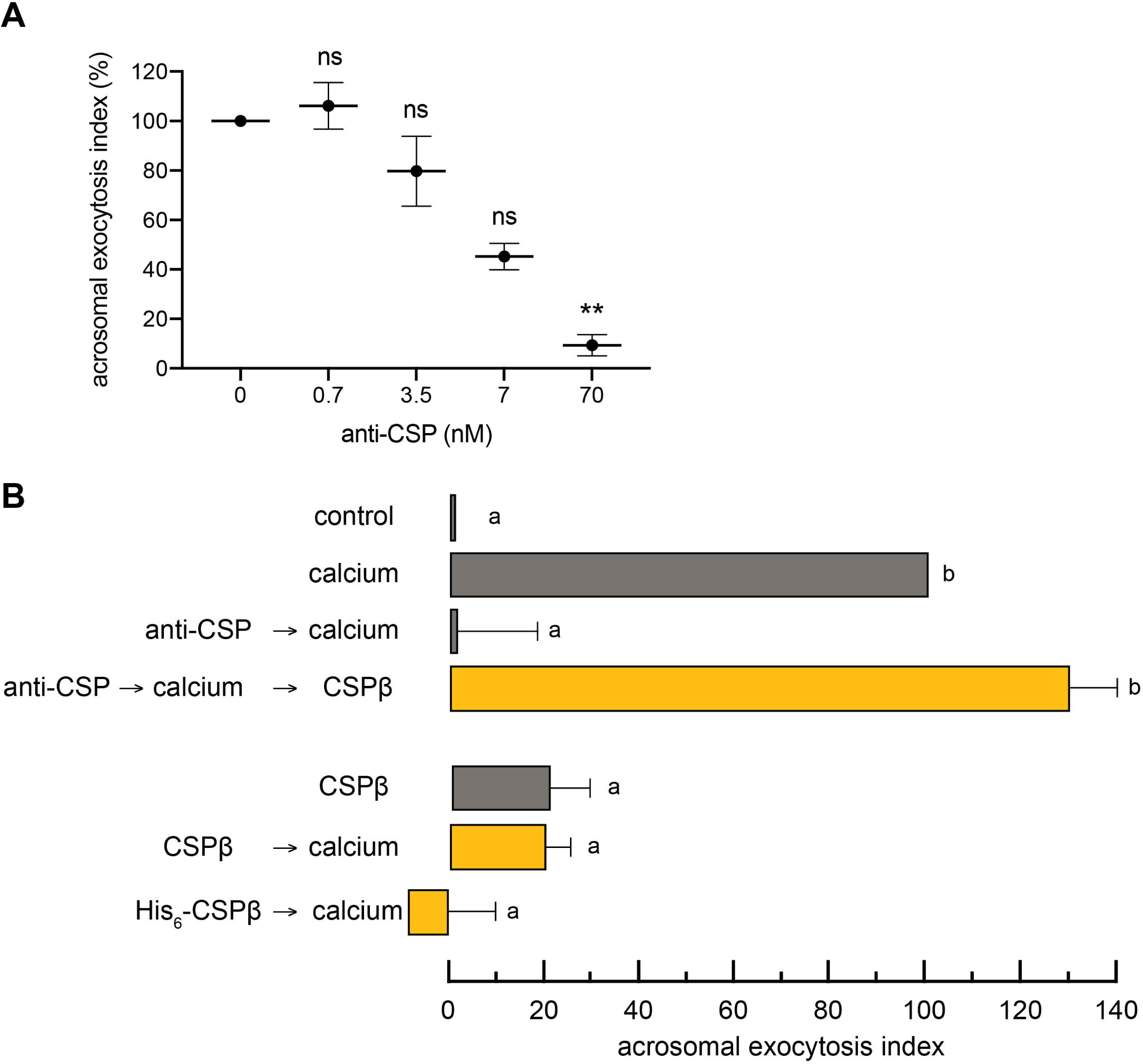
CSP is necessary for acrosomal exocytosis in human sperm. (**A**) SLO-permeabilized sperm were incubated for 15 min at 37° C with increasing concentrations of anti-CSP. Afterward, the sperm were incubated for 15 min at 37° C with 0,5 mM CaCl_2_ to initiate the acrosomal exocytosis. Asterisks indicate statistical significance (* * *p*_J<_J0.01), and ns indicates that statistical difference was nonsignificant (*p*_J>_J0.05) when compared with the acrosomal exocytosis index of 0 nM anti-CSP. (**B, top**) SLO-permeabilized sperm were incubated first with 70 nM anti-CSP for 15 min at 37° C and then were stimulated with 0.5 mM CaCl_2_ to induce secretion for another 15 min at 37° C. Finally, they were incubated with 140 nM recombinant GST-CSPβ to rescue exocytosis for 15 min at 37° C (yellow bar, anti-CSP→calcium→CSPβ). Controls (grey bars) included permeabilized sperm without stimulus (control), with 0,5 mM CaCl_2_ (calcium) and inhibition of calcium triggered exocytosis by 70 nM anti-CSP (anti-CSP→calcium). (**B, bottom**) SLO-permeabilized sperm were incubated with 140 nM GST-CSPβ (orange bar) or His_6_-CSPβ (yellow bar) for 15 min at 37° C and then were stimulated with 0.5 mM CaCl_2_ to induce secretion for another 15 min at 37° C. As control (grey bar) some aliquots were incubated with 140 nM GST-CSPβ alone (CSPβ). For all panels, acrosomal exocytosis was evaluated using FITC-PSA. Data were normalized as indicated in Materials and Methods. Plotted data represent the mean_J±_JSEM of at least three independent experiments. Different letters indicate statistical significance (*p*_J<_J0.01).

Different studies in other cellular models, show that diminished CSP expression or transient overexpression pointed to a decrease in regulated exocytosis (Brown *et al*., 1998; Zhang *et al*., 1998). On the other hand, stable overexpression of CSP augments exocytosis (Chamberlain and Burgoyne, 1998). As sequestration of CSP leads to acrosomal exocytosis inhibition in human sperm, we investigated the effect of the addition of recombinant CSPβ to SLO permeabilized human sperm. We tested two recombinant non-palmitoylated CSPβ (GST-CSPβ and His_6_-CSPβ) to prove that the tag did not affect the function of the CSP. Interestingly, when we added the recombinant CSPβ and triggered the exocytosis with calcium, it was abolished (Fig. 2B, bottom yellow bars) similarly to the antibody (Fig. 2B, top).

Our results indicated that blocking the effect of endogenous CSP, with anti-CSP or adding exogenous CSPβ, inhibited the acrosomal exocytosis. These data led us to wonder about the mechanism involved in this observation and explore the morphological changes in the sperm. Therefore, by transmission electron microscopy (TEM), we analysed the morphology of the acrosome and the membranes (plasma, inner and outer acrosomal) of the sperm head when we blocked CSP action, and consequently, acrosomal exocytosis was impaired. To do this, we performed assays with permeabilized sperm incubated with anti-CSP or recombinant CSPβ, and then stimulated with calcium. In these experiments, we included a negative control (not stimulated) and positive control (stimulated in the presence of 2-APB, an inhibitor of the acrosome reaction). Hence, swelling under the different conditions tested was always compared with the controls run in parallel. Fig. 3 presents the TEM assays performed. We showed representative images of the different patterns detected in the sperm head (Fig. 3A). In (a), a sperm with an intact acrosome with an electron-dense content and a flat outer acrosomal membrane-proximal and parallel to the plasma membrane, while in (b) and (c), a sperm with morphologically altered acrosomes. More specifically, we observed the following patterns: swelling of the acrosome granule, with the presence of waving in its membranes (b), and a phenotype similar to b but with apposed outer acrosomal and plasma membranes (c). Moreover, in (d), a reacted sperm that lacks the acrosome and where the inner acrosomal membrane becomes part of the limiting membrane of the cell.

**Figure 3.**
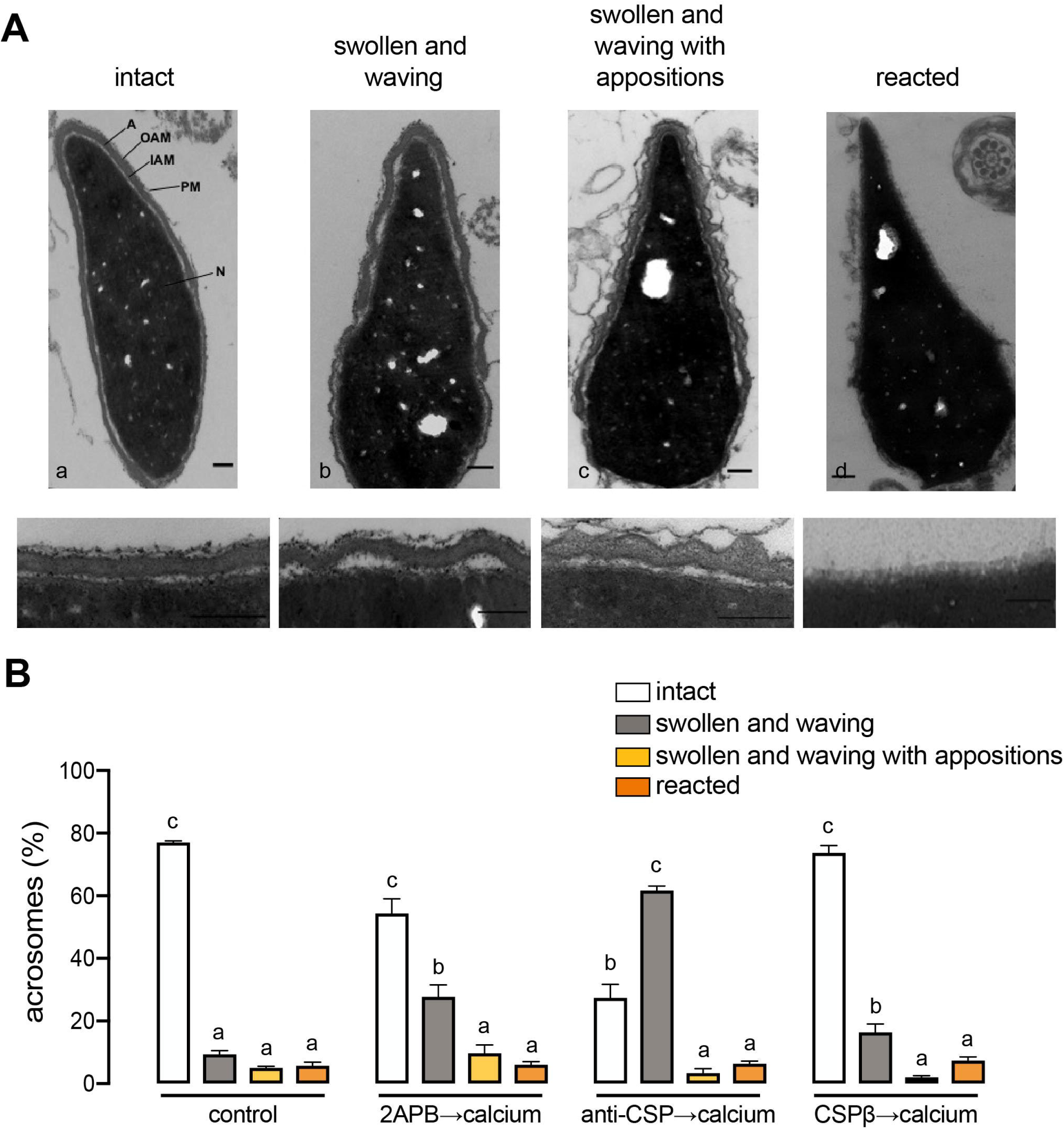
Morphological analysis by TEM after blocking the effect of endogenous CSP. Permeabilized sperm were incubated for 15 min at 37°C with 2-APB (200 µM) and then with anti-CSP (70 nM) or recombinant GST-CSPβ (400 nM). Subsequently, the cells were stimulated for 10 min at 37°C with 0.5 mM CaCl_2_. As a control, an aliquot was incubated in the absence of inhibitors and calcium (intact). Samples were fixed and processed for electron microscopy as described in Materials and Methods. (**A**) TEM micrographs illustrate different patterns of the acrosome, (a) intact, (b) swollen and waving, (c) swollen and waving with appositions, and (d) reacted. Down/bottom: higher magnification of the images showing details of the exocytic process. Scale bars 200 nm. (**B**) Percentage of acrosomes with the different morphological patterns observed, for each experimental condition: control, 2APB→calcium, anti-CSP→calcium, and CSPβ→calcium. Data were obtained from three independent experiments in which 200 cells were quantified for each condition. Different letters indicate statistical significance (p < 0.05, two-way ANOVA and Dunnett posthoc test). Abbreviations: A, acrosome, N, nucleus; PM, plasma membrane; OAM, outer acrosomal membrane; IAM, inner acrosomal membrane.

The TEM results showed that the alteration of the CSP function with the antibody mainly caused a swollen and waving pattern (Fig. 3B, anti-CSP→calcium “swollen and waving”, grey bar), and blocked the appositions between the membranes (only 3.3±1.4 %, Fig. 3B, swollen and waving with appositions, yellow bar). These data suggested that sequestering endogenous CSP would restrain the *trans*-SNARE complex association. On the other hand, the alteration of CSP activity with the recombinant CSPβ protein also showed morphologically altered acrosomes, but in this case, decreased the presence of swollen and waving acrosomes (Fig. 3B, 16.3±2.7% in CSPβ→calcium, grey bar; vs. 61.7±1.4% in anti-CSP→calcium, grey bar), and increased the percentage of intact acrosomes (Fig. 3B, 73.7 ± 2.3% in CSPβ→calcium, “intact” white bar; vs. 27.3±4.7% in the presence of anti-CSP→ calcium, “intact” white bar). This result suggests that the presence of recombinant CSPβ arrested exocytosis before acrosome swelling.

### CSP is required downstream NSF for the acrosomal exocytosis and is necessary for efficient stabilization of SNARE complexes in *trans* configuration

The results obtained from electron microscopy suggested a role of CSP in SNARE complex assembly which would stabilize membrane appositions. To test this hypothesis, we designed assays using reversible inhibitors. We know that the acrosome exocytosis is dependent on an extracellular calcium influx and internal store efflux (acrosome) into the cytosol sperm (Costello *et al*., 2009; Darszon *et al*., 2011). Thereby, we can reversibly stop the signaling cascade leading to the exocytosis using a membrane-permeant photolabile calcium chelator, NP-EGTA-AM, that sequesters intra-acrosomal calcium (De_Blas *et al*., 2002). To do this, we loaded the permeabilized sperm with NP-EGTA-AM in the dark and stimulated acrosomal exocytosis, allowing the signaling cascade to proceed until the acrosomal calcium was needed. Then, we released the caged calcium with UV light pulse resuming the exocytosis. In a control condition, when we added the anti-CSP before the inducer (calcium) to block the function of CSP, the exocytosis was prevented. However, incubation with anti-CSP after the inducer failed to inhibit acrosomal exocytosis (Fig. 4A top, yellow bar). Because recombinant CSPβ inhibited the acrosomal exocytosis (Fig. 2C), we used the NP-EGTA-AM assay to confirm if it performs in the same step as the endogenous one. As anticipated, recombinant CSPβ affected exocytosis before intra-acrosomal release (Fig. 4A bottom, yellow bar).

**Figure 4.**
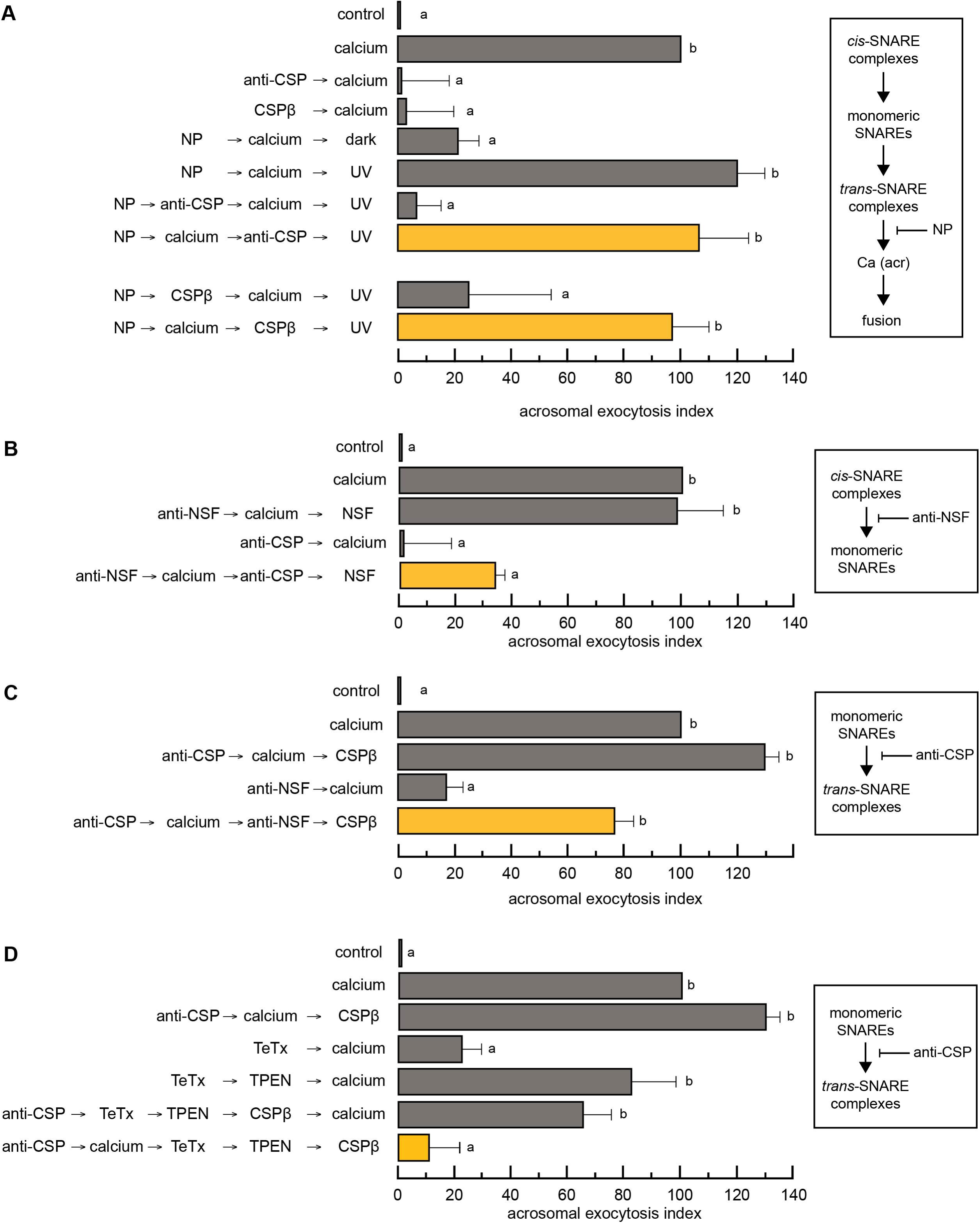
CSP is required downstream NSF for the acrosomal exocytosis and is necessary for SNARE complexes assembly in *trans* configuration. (**A**) SLO-permeabilized sperm were loaded with 10 µM NP-EGTA-AM (NP) in the dark for 10 min at 37° C to chelate intra-acrosomal calcium. Then the acrosomal exocytosis was stimulated with 0.5 mM calcium CaCl_2_ for another 15 min at 37° C, allowing the signaling cascade to proceed until the acrosomal calcium is needed. Next, sperm were treated with 70 nM anti-CSP (top) or 140 nM recombinant GST-CSPβ (bottom) for 15 min at 37° C. This procedure was carried out in the dark. We released the caged calcium with a UV light pulse resuming the exocytosis (NP→calcium→anti-CSP→UV, top yellow bar; NP→calcium→CSPβ→UV, bottom yellow bar). In a control condition, the anti-CSP or CSPβ were added for 15 min at 37° C before the inducer (calcium) to block the function of CSP, and then the exocytosis was triggered by 0.5 mM CaCl_2_ for 15 min at 37° C (NP→anti-CSP or CSPβ →calcium→UV). Several controls were conducted, some aliquots were incubated without CaCl_2_ (control), with CaCl_2_ in the absence of inhibitors (calcium), the inhibitory effect of NP-EGTA-AM in the dark (NP→calcium→dark), the recovery upon UV light pulse (NP→calcium→UV). (**B**) SLO-permeabilized sperm were treated with 1:200 anti-NSF for 15 min at 37° C before the inducer (calcium) to block the function of NSF. Then the exocytosis was triggered by 0.5 mM CaCl_2_ for 15 min at 37° C, and next 70 nM anti-CSP for another 15 min at 37° C. Finally, the blockage of anti-NSF was rescued by an additional 15 min at 37° C in the presence of 60 nM NSF (anti-NSF→calcium→anti-CSP→NSF, yellow bar). Several controls were conducted, some aliquots were incubated without CaCl_2_ (control), with CaCl_2_ (calcium), the recovery with 60 nM recombinant NSF (anti-NSF→calcium→NSF), and the inhibitory effect of 70 nM anti-CSP (anti-CSP→calcium). (**C**) SLO-permeabilized sperm were treated with 70 nM anti-CSP for 15 min at 37° C. The acrosomal exocytosis was initiated by adding 0.5 mM CaCl_2_ and incubating for 15 min at 37° C. Then, 1:200 anti-NSF was added for another 15 min at 37° C, and finally, the blockage was rescued by an additional 15 min at 37° C in the presence of 140 nM GST-CSPβ (anti-CSP→calcium→anti-NSF→CSPβ, yellow bar). Several controls were conducted, some aliquots were incubated without CaCl_2_ (control), with CaCl_2_ (calcium), rescue with 140 nM recombinant CSPβ (anti-CSP→calcium→CSPβ), and the inhibitory effect of 1:200 anti-NSF (anti-NSF→calcium). (**D**) SLO-permeabilized sperm were incubated for 10 min at 37°C with 70 nM anti-CSP, and then stimulated with 0.5 mM CaCl_2_ for 10 min at 37°C to initiate exocytosis. Then, 100 nM TeTx was added and left to act for 10 min at 37° C, next the sperm were treated with 2.5 µM TPEN for another 10 min at 37° C, and finally incubated with 140 nM recombinant CSPβ for 10 min at 37° C to resume exocytosis (anti-CSP→calcium→TeTx→TPEN→CSPβ, yellow bar). As control (gray bars), cells were incubated without any treatment (control), with 0.5 mM CaCl_2_ (calcium), the exocytosis rescue with 140 nM recombinant CSPβ (anti-CSP→calcium→CSPβ), the inhibitory effect of 100 nM TeTx (TeTx→calcium), impairing of toxin cleavage by 2.5 µM TPEN (TeTx→TPEN→calcium), and recovery of anti-CSP blockage before the addition of CaCl_2_ (anti-CSP→TeTx→TPEN→CSPβ→calcium). Acrosomal exocytosis was evaluated using FITC-PSA. Data were normalized as indicated in Materials and Methods. Plotted data represent the mean_J±_JSEM. Different letters indicate statistical significance (*p* <_J0.01) from at least three independent experiments.

Previous research of our lab found that NSF is required for acrosomal exocytosis (Tomes *et al*., 2005), NSF/α-SNAP disassembled *cis* complexes into monomeric SNAREs. Based on this finding and our results, we propose that CSP participate in SNARE assembly in *trans* configuration, so we explored if CSP is performing its action after NSF. Our model of permeabilized sperm allows us to pause triggered exocytosis and establish the exact moment where the protein is needed in the signaling cascade. We used an approach called “reversible pair,” which consists of an exocytosis blocker that sequesters a protein essential for fusion and the recombinant protein that reverses the blockade imposed by the antibody (Ruete *et al*., 2014). First, we used anti-NSF/recombinant NSF reversible pair (Fig. S1). We predicted that adding the anti-NSF and calcium before anti-CSP and then recovering with recombinant NSF will block the exocytosis. As we expected, this sequence inhibited the acrosomal exocytosis (Fig. 4B, anti-NSF→calcium→anti-CSP→NSF, yellow bar). To confirm our results, we performed a similar approach with a second reversible pair anti-CSP/recombinant CSPβ. We anticipated that if NSF is required before CSP, incubating with anti-CSP before calcium, then with anti-NSF, and finally, adding CSPβ will conduct to acrosomal exocytosis. According to our expectations, this was the case (Fig. 4C, anti-CSP→calcium→anti-NSF→CSPβ, yellow bar). Both results confirmed that CSP worked after NSF.

During capacitation and in resting sperm, SNAREs are resistant to neurotoxin because they are engaged in *cis* complexes (Tomes *et al*., 2002; De_Blas *et al*., 2005). As mentioned above, the *cis* complexes are disassembled by the addition of NSF/α-SNAP (monomeric SNAREs), becoming sensitive to toxin cleavage before the intra-acrosomal calcium is released (De_Blas *et al*., 2005). In the final steps of membrane fusion, SNAREs assemble in *trans* complexes. Since we showed that CSP acts after NSF, and based on electron microscopy results that the sequestering of CSP avoided the presence of membrane appositions (Fig. 3B), we wondered if CSP has a role in SNARE assembly in *trans* configuration. To determine the effect of CSP in the state of SNARE assembly, we carried out exocytosis assays using the light chain of tetanus toxin (TeTx), a protease specific for monomeric synaptobrevin-2. We can use the TeTx to monitor SNARE configuration. Assembled *trans*-SNARE complexes are resistant to TeTx because synaptobrevin-2 is protected from proteolytic cleavage. As a control, we treated permeabilized sperm in the absence or presence of the light chain of TeTx in the absence or presence of calcium. When indicated, we treated the sperm with TPEN, a Zn^2+^ chelator, to stop the toxin effect (De_Blas *et al*., 2005). We incubated permeabilized sperm with anti-CSP and then with calcium to start exocytosis. Then we added TeTx and let the toxin act, next we treated the sperm with TPEN, and finally, we incubated it with the recombinant CSPβ to resume exocytosis. Under these conditions, calcium did not accomplish exocytosis, showing that anti-CSP prevented *trans*-SNARE complex assembly. (Fig. 4C bottom, yellow bar).

## DISCUSSION

Even though CSP was discovered more than 20 years ago, most published research focuses on CSP’s role in neurodegenerative diseases (reviewed in (Gundersen, 2020)). Nevertheless, little is known about its role in acrosomal exocytosis and its fertility implications. A portion of men has unexplained male infertility, despite having normal semen analysis. In searching for possible players involved in cellular sperm dysfunction, the present study focused on the role of CSP in acrosomal exocytosis in human sperm cells, and this event is crucial for oocyte fertilization. Here, we report that CSP is present in human sperm cells and has a role in stabilizing *trans*-SNARE complexes during acrosomal exocytosis.

DnaJC5β expression predominantly in testis, and single-cell RNA cluster analysis of CSPβ in late and early spermatids reinforce the idea that this protein could be implicated in sperm physiology. The molecular weight of endogenous CSP corresponds to a full palmitoylated version of CSP (Coppola and Gundersen, 1996), demonstrating that this protein is palmitoylated in human sperm. Further studies are needed to understand each CSP isoform’s relative contribution to the human sperm exocytosis (the antibody recognizes CSPα and β). Nevertheless, our data propose that the main CSP isoform detected in our experimental setting is CSPβ, given their preferential expression in testis. CSP localization in the head sperm is consistent with a protein involved in the acrosome reaction. In neurons, CSP is attached to the synaptic vesicle membrane through palmitoylation (Zinsmaier *et al*., 1990; Ohyama *et al*., 2007). We observed that CSP localizes to particulate fraction and partitions into the Triton X-114 detergent phase, suggesting that endogenous CSP is attached to membranes and is palmitoylated in the human sperm extract.

The experiments sequestering endogenous CSP with a specific antibody inhibited calcium triggered exocytosis, indicating that the presence of CSP is necessary for acrosomal exocytosis. As we proposed in a recent work (Ruete *et al*., 2014), the rescue of this inhibition by adding recombinant CSPβ could be explained by the sequestration of endogenous CSP by the antibody and restricting its function. The addition of the recombinant CSPβ displaces the antibody from the endogenous CSP allowing the exocytosis to continue and again confirming that anti-CSP recognized CSPβ. Consistent with our hypothesis, we conclude that these results sustain the notion that CSP is required for acrosomal exocytosis and confirm the critical role of CSP in this process.

We showed that in sperm, endogenous CSP is palmitoylated and anchored on the membrane, probably through palmitoylation by the cysteines located in the cysteine string domain. We used a recombinant CSPβ produced in bacteria that lacks this posttranslational modification, and so, the properties expected of a membrane-bound protein. Recombinant CSPβ inhibited the acrosomal exocytosis triggered by calcium, so we concluded that a mislocalization of recombinant CSPβ would be possible, avoiding the interaction of CSPβ and the sperm membranes. Tobaben *et al*. (Tobaben *et al*., 2001) reported a trimeric protein complex composed of CSP, Hsc70 (heat shock cognate 70), and SGT (small glutamine-rich tetratricopeptide repeat protein) that functions as a synaptic chaperone machine in mice neurons. We detected the presence of these proteins in human sperm (unpublished data). We infer that a reasonable explanation for the exocytosis inhibition in the presence of CSPβ might be that exogenous non-palmitoylated mislocalized CSPβ, could sequester endogenous Hsc70 or SGTA. Sequestering one of these proteins participating in the complex would avoid the correct subcellular localization and function (Rodríguez *et al*., 2012).

Acrosome swelling is crucial for the close up of outer acrosomal and plasma membranes leading to acrosomal exocytosis (Zanetti and Mayorga, 2009). They propose that *trans*-SNARE complexes are assembled in membrane apposition regions and prelude the fusion process. Many factors interact with the *trans*-SNARE complex to regulate its assembly, but it is still unclear whether these interactions occur (Rizo and Rosenmund, 2008). The electron microscopy images of “swollen and waving acrosomes” in the presence of anti-CSP or recombinant CSPβ and the absence of membrane appositions agreed with our prior findings that CSP is necessary for exocytosis and suggests its participation in the *trans*-SNARE complex assembly. Interestingly, the increase in the “intact” pattern in the presence of recombinant CSPβ suggests a different mechanism for the inhibition of exocytosis. However, further studies are needed to confirm this hypothesis.

We showed that endogenous CSP and recombinant CSPβ participate in the signaling cascade before intra-acrosomal calcium release, and these results are consistent with CSP is required to stabilize *trans*-SNARE complexes. Previous works from our lab indicated that *cis*-SNARE are disassembled by NSF/α-SNAP (De_Blas *et al*., 2005; Zarelli *et al*., 2009; Rodríguez *et al*., 2011). We revealed that CSP performs its role downstream of NSF in the acrosomal exocytosis under these experimental conditions. Our results are consonant with studies in chromaffin cells in which there is also a one-shot fusion event. Graham and Burgoyne (Graham and Burgoyne, 2000) demonstrate that α-SNAP acts in early fusion steps, and CSP plays close to the fusion process. More work needs to be done to know the CSP clients in human sperm and the possible implications of its impairment in fertility.

In our sperm model, we can reversible inhibit the calcium efflux from the acrosome and block the exocytosis before the *trans-SNARE* complex assembles. The inhibition of acrosomal exocytosis when sequestering CSP showed that synaptobrevin2 becomes sensitive to TeTx. This observation is consistent with our previous findings, where there are no membrane appositions because no *trans-*complexes were formed in the presence of anti-CSP. These results show that CSP stabilizes the *trans-*configuration (Fig. 5), supporting the role of CSP in promoting the preservation of SNARE machinery and highlighting the importance of studying this protein in idiopathic infertility cases. A neglected area in acrosomal exocytosis is understanding the protein interactions between CSP and SNAREs and accessory proteins to the SNARE assembly process. Previous work in several cellular models shows that CSP interacts with synaptotagmin 9 and the SNARE proteins SNAP-25, synaptobrevin-2, and syntaxin 1 (reviewed in (Gundersen, 2020)). Additional studies are needed to elucidate the precise mechanism by which CSPβ fine-tune acrosomal exocytosis in human sperm.

**Figure 5.**
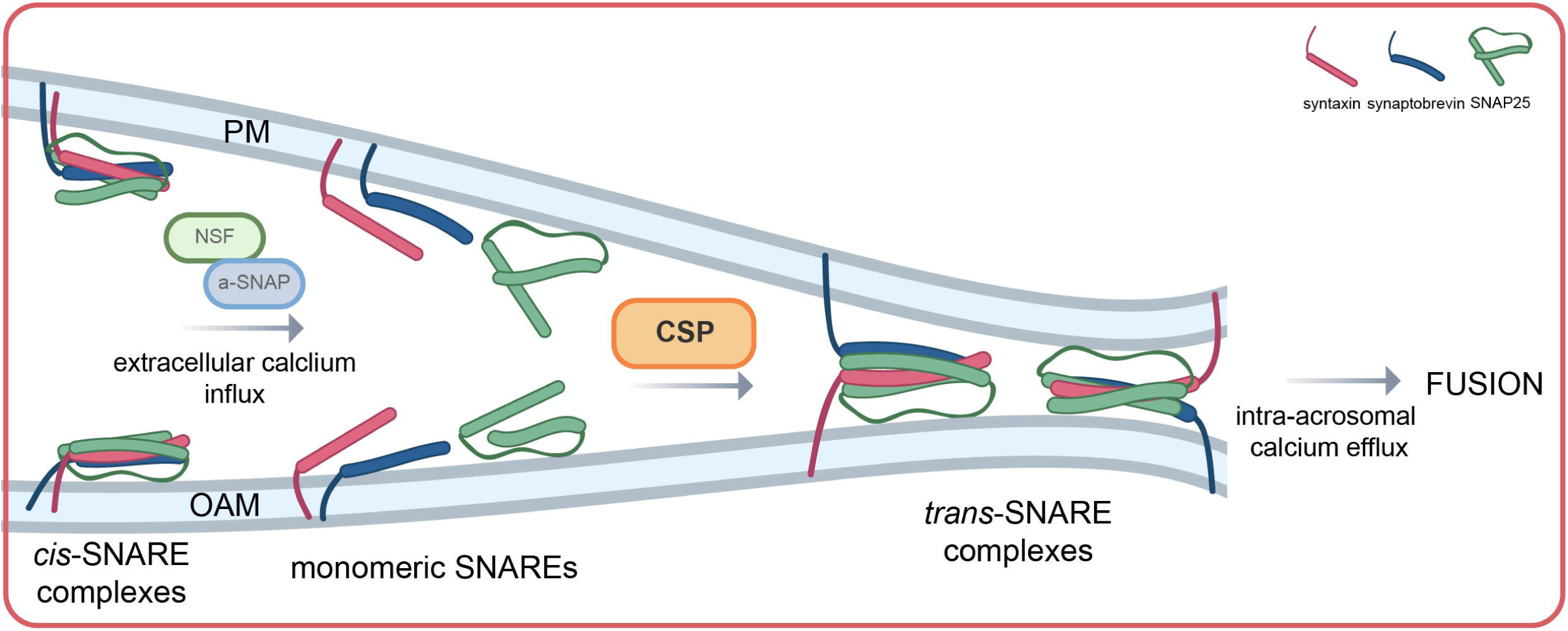
Working model for CSP during acrosomal exocytosis. In resting sperm, SNAREs are assembled in *cis*-complexes. Following extracellular calcium influx, NSF/α-SNAP disassemble *cis* configuration into monomeric SNAREs. Now SNAREs are stabilized in *trans* configuration by the action of CSP. An intra-acrosomal calcium efflux led to membranes fusion. PM, plasma membrane; OAM, outer acrosomal membrane.

In closing, CSP controls one of the key processes for male fertility. Understanding its role is critical in identifying new biomarkers and generating new rational-based approaches to treating male infertility.

## MATERIAL AND METHODS

### Reagents

We obtained recombinant streptolysin O (SLO) from Dr. Bhakdi (University of Mainz, Mainz, Germany). Spermatozoa were cultured in Human Tubal Fluid media (as formulated by Irvine Scientific, Santa Ana, CA, USA, HTF) supplemented, when indicated, with 0.5% bovine serum albumin (BSA). Human CSPβ fused to His_6_ in pET28a was from GenScript (NJ, USA). The rabbit polyclonal anti-CSP antibody (affinity-purified with the immunogen directed towards the amino acids 182-198 of rat CSP), the rabbit polyclonal anti-NSF (antiserum), and monoclonal mouse anti-alpha-tubulin (purified IgG) were from Synaptic Systems (Göttingen, Germany). The rabbit polyclonal anti-GST (purified IgG) antibody was from Novus Biologicals, LLC (Centennial, CO, USA). Horseradish peroxidase anti mouse, anti-rabbit, and Cy™ 3-conjugated goat anti-rabbit IgGs (H+L) were from Jackson ImmunoResearch (West Grove, PA). We obtained the synaptosomal preparation from Dr. V. Gonzalez Polo and Dr. S. Patterson (University of Cuyo, Mendoza, Argentina). 2-aminoethoxydiphenyl borate (2-APB) from Calbiochem was purchased from Merck Quí mica Argentina SAIC (Buenos Aires, Argentina). O-nitrophenyl EGTA-acetoxymethyl ester (NP-EGTA-AM) was purchased from Life Technologies (Buenos Aires, Argentina). N, N, N’, N’-tetrakis (2-pyridymethyl) ethylenediamine (TPEN) was from Molecular Probes (Waltham, MA, USA). *Pisum sativum agglutinin* (PSA) lectin labelled with fluorescein isothiocyanate (FITC-PSA), paraformaldehyde, poly-L-lysine, bovine serum albumin (BSA), and Tannic acid were acquired from Sigma-Aldrich™ (Buenos Aires, Argentina). Isopropyl-D-1-thiogalactopyranoside (IPTG) was purchased from ICN (Eurolab SA, Buenos Aires, Argentina). All electron microscopy supplies were from Pelco (Ted Pella Inc. CA, USA). All other chemical reagents were of analytical grade and were purchased from ICN, Sigma-Aldrich™, Genbiotech, or Tecnolab (Buenos Aires, Argentina).

### Recombinant proteins

We use the following constructs for protein expression in *Escherichia coli* BL21(DE3) cells: full-length human CSPβ fused to GST in pGEX-2T was kindly provided by Dr. P Scotti (Laboratoire de Chimie et Biologie des Membranes et Nano-objets, Université de Bordeaux, France), human CSPβ fused to His_6_ in pET28a was from GenScript (NJ, USA), full-length human NSF fused to His_6_ in pET28 was a kind gift from Dr. D. Fasshauer (Department of Fundamental Neurosciences, Université de Lausanne, Switzerland), and light chain TeTx fused to His_6_ in pQE3 was generously provided by Dr. T. Binz (Institut für Zellbiochemie, Medizinische Hochschule Hannover, Germany).

GST-full-length CSPβ expression was induced with 1 mM isopropyl-β-D-thio-galactoside (IPTG) for 4 h at 37° C. Purification was done using Glutathione Sepharose 4B (Amersham) in PBS pH 8, 5 mM β-mercaptoethanol, 10 mM MgCl_2_, 1 mM ATP, 10 μg/ml DNAse and 1% Triton X-100 followed by elution in 50 mM Tris HCl pH 8, 500 mM NaCl and 20 mM glutathione.

Expression of His_6_ proteins were performed as previously reported (Ruete *et al*., 2019) with the modifications described below. His_6_-full-length NSF was induced with 1 mM IPTG for 3 h at 37° C. His_6_-light chain TeTx was induced with 0.25 mM IPTG for 3 h at 37° C. Purification of His_6_-full-length NSF was done using Ni-NTA resin according to Qiagen’s instructions except all buffers contained 20 mM Tris HCl pH 7.4 (instead of 50 mM phosphate pH 8), 200 mM NaCl, 0.5 mM ATP, 5 mM MgCl2, and 2 mM β-mercaptoethanol followed by elution in 20 mM Tris HCl pH 7.4, 200 mM NaCl, 0.5 mM ATP, 5 mM MgCl_2_, 2 mM β-mercaptoethanol and 250 mM imidazole. Purification of His_6_-light chain TeTx was done using Ni-NTA resin (Qiagen) in 50 mM Tris HCl pH 7.4, 500 mM NaCl and 50 mM imidazole followed by elution in 50 mM Tris HCl pH 7.4, 300 mM NaCl and 350 mM imidazole.

Recombinant protein concentration was quantified through BCA protein assay kit (Thermo Fisher Scientific, Buenos Aires, Argentina) on a BioRad 3350 Microplate Reader using BSA as standard, or from intensities of the bands in Coomassie blue-stained, sodium dodecyl sulfate-polyacrylamide electrophoresis (SDS-PAGE) gels.

### Human sperm samples preparation

Our research followed ethical principles outlined in the Declaration of Helsinki. All experimental procedures for the collection and manipulation of human sperm samples were approved by the Bioethical Committee of the Medical School (Comité de Bioética de la Facultad de Ciencias Médicas de la Universidad Nacional de Cuyo, EXP-CUY: 25685/2016). Human semen samples were obtained from healthy volunteer donors (age range 21-45). An informed consent form was signed by donors.

Semen samples were liquefied for 30-60 min at 37 °C. The highly motile cells were separated from the seminal plasma by a swim-up protocol incubating for 1h in HTF, 37° C and 5%CO_2_/95% air conditions. Briefly, sperm cells were incubated at a concentration of 10^7^/ml during 2 h under capacitating conditions (HTF supplemented with 0.5% BSA, 37 °C, 5% CO_2_/95% air). Then, the capacitated sperm cells were washed with PBS and permeabilized in cold PBS containing 3 U/ml SLO for 15 min at 4° C. Next, the sperm were resuspended in a sucrose buffer containing 250 mM sucrose, 0.5 mM EGTA, 20 mM Hepes-K, pH 7.0, and 2 mM DTT. Then samples were prepared for acrosomal exocytosis assays, indirect immunofluorescence, and Transmission Electron Microscopy.

### Acrosomal exocytosis assays

Capacitated and permeabilized sperm were treated sequentially with inhibitors and stimulants according to the assay, as indicated in the figure captions and incubated for 10-15 min at 37° C after each addition. When indicated, samples were loaded with NP-EGTA-AM (a photosensitive intracellular calcium chelator) in the dark for 10 min at 37° C without calcium. Then the cells were treated in the presence of inhibitors and calcium. After the incubations, the sperm were exposed twice to UV flash (1 min each time at 37° C). Finally, the samples were processed for acrosomal exocytosis evaluation, as described in (Ruete *et al*., 2019). The acrosomal status was determined by using fluorescein isothiocyanate-conjugated *Pisum sativum* agglutinin (FITC-PSA) staining according to (Mendoza *et al*., 1992).

### Indirect immunofluorescence

Sperm were capacitated for 2 h and fixed in 2% paraformaldehyde/PBS for 15 min at room temperature. Then they were resuspended in 100 mM glycine/PBS to stop the fixing. After, the sperm were attached to 12 mm round coverslips treated with 0,005% poly-L-lysine (w/v) in distilled water for 30 min at room temperature in a moisturized chamber. The plasma membrane was permeabilized with 0.1% Triton X-100 in PBS for 10 min at room temperature and washed three times with 0.1% polyvinylpyrrolidone (PVP-40) in PBS (PBS/PVP). To block nonspecific staining in the sperm, they were treated for 1 h at 37° C with 5% BSA in PBS/PVP. Next, the sperm were incubated for 1 h at 37° C with anti-CSP (10 µg/ml) diluted in 3% BSA in PBS/PVP. Later, they were washed three times with PBS/PVP and treated for 1 h at 37° C with Cy™3-conjugated anti-rabbit IgG (2.5 µg/ml in 1% PBS/PVP). Then, coverslips were washed three times with PBS/PVP, and the sperm plasma membrane was permeabilized for 20 sec with ice-cold methanol. Acrosomes were stained for 40 min with FITC-PSA (25 µg/ml in PBS) and washed for 20 min with distilled water. Finally, the coverslips were mounted with Mowiol^®^ 4-88 in PBS supplemented with 2 µm Hoechst 33342. The samples were examined with an Eclipse TE2000 Nikon microscope equipped with a Plan Apo 60x/I.40 oil objective. The images were taken with a Hamamatsu digital C4742-95 camera operated with Metamorph 6.1 software (Universal Imaging, Bedford Hills, NY, USA). We counted at least 200 cells per condition.

### Subcellular fractionation

The protocol described by (Bohring and Krause, 1999) and modified by (Tomes *et al*., 2005) was followed. The capacitated sperm (10 × 10^6^ cells/ml) were diluted 1:9 in a hypoosmotic buffer (Jeyendran *et al*., 1984) and incubated for 2 h at 37° C. The samples were sonicated three times for 15 min at 40 Hz on ice, and centrifuged at 10,600 x g for 15 min at 4° C. An additional centrifugation at 20,800 x g was done to remove cell debris. Finally, a 208,000 x g centrifugation for 2 h at 4° C separated pellets (particulate fraction) from supernatants (soluble fraction).

For phase separation in Triton X-114, sperm were treated following standard procedures (Bordier, 1981) and modified by (Bustos *et al*., 2012).

### SDS-PAGE and immunoblotting

Equal amounts of protein were resolved on 10% SDS-PAGE and blotted onto nitrocellulose membranes (GE Healthcare). Immunoblots were blocked with 5% fat-free milk in PBS containing 0.1% Tween-20 for 1h at room temperature. Then, membranes were probed with anti-CSP (1 μg/ml) or anti-tubulin (1 μg/ml) at 4° C overnight. The following day, blots were incubated with HRP-conjugated anti-rabbit IgG (for anti-CSP) or goat anti-mouse IgG (for anti-tubulin) (0.1 μg/ml in PBS) for 1 h at room temperature. Immunoreactive proteins were detected with a chemiluminescence system (Kallium Technologies, Buenos Aires, Argentina) using a Luminescent Image Analyzer LAS-4000.

### Transmission Electron Microscopy assays of the coincubation of anti-CSP and GST-CSP with sperm

Human spermatozoa was processed as described earlier (Zanetti and Mayorga, 2009). Then, cells were fixed at room temperature (RT) with 2.5% (w/v) glutaraldehyde in PBS, pH 7.4 for 1 h. Fixed samples were washed twice in PBS and post-fixed in 1% (w/v) osmium tetroxide-PBS for 1 h at room temperature, and dehydrated with increasing concentrations of cold acetone. Cells were infiltrated at room temperature in 1:1 acetone:Spurr for 2 h, and finally embedded in fresh pure resin overnight at RT. Samples were cured 24 h at 70°C. A diamond knife (Diatome) was used to cut thin sections (80 nm) on a Leica Ultracut R ultra-microtome. Then, the samples were stained with uranyl acetate/lead citrate. TEM grids were photographed with a Zeiss 900 electron microscope at 80 kV. Representative images were selected for the manuscript. We included negative (sperm not stimulated) and positive (stimulated with 0.5 mM CaCl_2_ in the presence of 200 μM 2-APB) controls in all experiments.

### Statistical analysis

Prism 8 software was used for statistical analysis (GraphPad, La Jolla, CA, USA). Two-way ANOVA and Tukey post-test were used for multiple comparisons. Student’s t-test was used for unpaired data. Differences were considered statistically significant at the P-values < 0.05. For transmission electron microscopy Two-way ANOVA shows statistically significant difference between the different patterns, p < 0.05 (Dunnett’s t-test).

## Acknowledgments

The authors greatly acknowledge Dr. D. Croci for critical reading and helpful comments for manuscript revision, and to R. Militello, E. Bocanegra, N. Domizio and J. Ibañ ez for technical assistance.

## Data Availability

The data underlying this article will be shared on reasonable request to the corresponding author.

